# Analysis of pluripotency transcription factor interactions reveals preferential binding of NANOG to SOX2 rather than NANOG or OCT4

**DOI:** 10.1101/2020.06.24.169185

**Authors:** Tapan Kumar Mistri, David Kelly, John Mak, Douglas Colby, Nicholas Mullin, Ian Chambers

## Abstract

The pluripotency transcription factors (TFs) Nanog, Sox2, and Oct4 are at the centre of the gene regulatory network that controls cell identity in embryonic stem (ES) cells. However, the mechanisms by which these factors control cell fate, and their interactions with one another are not fully understood. Here we combine biophysical and novel biochemical assays to assess how these factors interact with each other quantitatively. A confocal microscopy method to detect binding of a target protein to a fluorescently labelled partner (coimmunoprecipitated bead imaging microscopy [CBIM]) is presented and used to demonstrate homotypic binding of Nanog and heterotypic binding between Nanog and Sox2 and between Nanog and Oct4. Using fluorescence correlation spectroscopy we show that in solution, Nanog but not Oct4 or Sox2 can form homomultimers. However, both Sox2 and Oct4 can form heterotypic multimers with Nanog in a manner that depends on the presence of tryptophan residues within the Nanog tryptophan repeat. Fluorescence Cross Correlation Spectroscopy shows the affinity of Nanog for dimer formation is in the order Sox2 > Nanog > Oct4. Importantly, live cell analysis demonstrates the existence of Nanog homomultimers *in vivo*. Together these findings extend understanding of the molecular interactions occurring between these central mediators of the pluripotency gene regulatory network at the single-molecule level.

## INTRODUCTION

ES cells originate from the inner cell mass (ICM) of the preimplantation mammalian embryo (Evans & Kaufman 1981) and are defined by their capacity for indefinite self-renewal in culture and their ability to differentiate into all three embryonic germ layers (Smith 2001). A network of core transcription factors (TFs) including Sox2, Oct4, and Nanog controls ES cell self-renewal efficiency (Boyer et al 2005, Chambers 2005). In particular, Nanog can enhance ES cellself-renewal to the point of cytokine independence, when overexpressed (Chambers et al 2003). Genetic knock-out of *Nanog* results in preimplantation lethality due to a failure to properly specify pluripotent cell identity (Mitsui et al 2003, Silva et al 2009). In ES cells, conditional deletion of *Nanog* increases the propensity of cells for differentiation, but does not eliminate pluripotency (Chambers et al 2007). These observations led to the hypothesis that Nanog functions as a rheostat with the level of Nanog positively correlated to ES cell self-renewal (Chambers et al 2003, Chambers et al 2007).

Previous studies have demonstrated that Nanog functions as a dimer (Mullin et al 2008, Wang et al 2008), with dimerisation occuring through a low complexity domain referred to as thetryptophan repeat (WR), due to the presence of a tryptophan at every fifth position (Mullin et al 2008). The dimerisation of Nanog may also be affected by additional Nanog partner proteins. While the residues involved in Nanog homodimerisation and heterodimerisation with Sox2 have been determined (Gagliardi et al 2013, Mullin et al 2017, Mullin et al 2008), how Nanog interacts with additional partners remains to be shown. For example, while Nanog has been reported to bind Oct4 (Wang et al 2008), other studies did not report an interaction between Nanog and Oct4 (Gagliardi et al 2013, van den Berg et al 2010). Therefore, the differential binding specificity of Nanog with specific partners requires further exploration.

In the current study, we generated and functionally characterised fluorescent GFP and mCherry tagged fusions with Nanog to investigate the molecular interactions of Nanog with Sox2 and Oct4 using FCS and FCCS (Chen et al 2012, Mistri et al 2015). We used single-molecule, sensitive techniques including fluorescence correlation spectroscopy (FCS) and cross-correlation spectroscopy (FCCS) (Macháň & Wohland 2014) to help advance understanding of how TFs interact with their binding partners both *in vitro* and under physiological conditions in live cells. In addition, we used a novel technique termed <u>c</u>oimmunoprecipitated <u>b</u>ead imaging <u>s</u>pectroscopy (CBIM) to directly visualise DNA-protein interactions. Our results reveal that Nanog has unique molecular binding dynamics among master regulators like Sox2 and Oct4.

## MATERIALS AND METHODS

### Cell culture and transfection

Chinese hamster ovary (CHO) cells (K1 line, ATCC # CRL**-**61) were cultured in Glasgow’s modified Eagle’s medium (GMEM, Life Technologies), 10% fetal bovine serum (GIBCO), 1x nonessential amino acids, 2 mM L**-**glutamine/pyruvate, 112 µM β-mercaptoethanol at 37 ºC, 7% CO_2_ and 95 % humidity. Transfection of plasmids was performed using Lipofectamine 3000 (Invitrogen) according to the manufacturer’s protocol. 10 µg of plasmid was transfected into CHO cells on 75cm^2^ plates. The cells were collected after 24 hours for nuclear lysate preparation. Mouse E14/T cells were cultured on gelatin-coated dishes in GMEM/β**-**mercaptoethanol/10% FCS/LIF as described earlier (Chambers et al 2003)

### Nuclear lysate preparation

Nuclear lysates were prepared from transfected CHO cells 24 hours post-transfection as described previously (Hutchins et al 2013, Mistri et al 2015).

### Co-immunoprecipitated bead imaging microscopy (CBIM)

Coimmunoprecipitation (Co-IP) transfections were performed with plasmids encoding (FLAG)_3_-tagged interaction partners or partners tagged with a fluorescent proteins using Lipofectamine 3000, (Life technologies) according to the manufacturer’s protocol. The nuclear lysate was prepared 24 hours post**-** transfection of 6 million cells in a final volume of 250 µl. The Co-IP was performed as described (Gagliardi et al 2013). After Co-IP, 30 µl of binding buffer was added to protein G sepharose beads. 10 µl of the bead solution was used for fluorescence detection by confocal microscopy. GFP and mCherry were imaged at a laser intensity of 31% with 488 nm and 561 nm lasers respectively. High laser power was used to eliminate background noise. For protein-DNA interactions, CoIP beads were incubated with Cy5-tagged DNA oligos, at a final concentration of 50 nM in 100 µl of binding buffer for 1 hour at 4°C with rotation. The beads were washed with 1ml PBS buffer 5 times before imaging. Cy5 tagged beads were imaged with a 633 nm laser at 31% of laser intensity.

### Fluorescence correlation and cross-correlation spectroscopy (FCS/FCCS)-

FCS was performed using a laser scanning confocal microscope (Leica SP5, Leica Microsystems UK) with a water immersion objective (63×, NA1.2, Leica) coupled to an in-built FCS module. A dichroic mirror (560DRLP) and bandpass emission filters (500-550nm for Alexa 488/GFP and 607-683 nm for Atto 565/mCherry, (Chroma Technology, VT) were used to separate the excitation light coming from the fluorescence emission. A standard calibration approach was used to determine the absolute concentration of the fusion proteins in the crude nuclear lysate (Mistri et al 2018, Mistri et al 2015). Appropriately chosen dyes with known diffusion coefficients were employed to determine the confocal volume accurately. To measure the concentration of GFP tagged proteins, calibration was performed with 2 nM of Alexa488 (diffusion constant, D = 3.80× 10^−6^ cm^2^s^-1^) (Ramadurai et al 2009) triggered by a 488 nm laser line at 30 µW (15% laser intensity). For mCherry-Sox2, the calibration was performed using a 561 nm laser line at 30 µW (10% laser power) using 2 nM of Atto565 (diffusion constant, D = 2.59 × 10^−6^ cm^2^s^-1^) (Pan et al 2006). For live-cell studies, GFP and mCherry localised to the nucleus were excited at 10 µW by 488 nm and 561 nm lasers respectively. Correlations in fluorescence fluctuations were computed online by a hardware correlator and were fitted using a theoretical model for free diffusion in solution and a triplet using Vista 3.6 (ISS, Inc Champaign IL) or using ain-house fitting module based on Igor Pro 6.0 as described previously (Wohland et al 1999). Absolute concentrations were calculated in nM.

## RESULTS

To assess the molecular interactions between pluripotency transcription factors (TFs), fluorescent protein fusions were prepared. Sox2 and Oct4 fluorescent protein fusions have previously been shown to be functional (Mistri et al 2015). For Nanog, fusions to mCherry or eGFP were prepared and their functionality tested following transfection of E14/T ESCs. This shows that both mCherry-Nanog and Nanog-GFP retain the defining capacity of Nanog to drive LIF-independent ESC self-renewal (Fig. S1).

### A bead-based assay for detecting protein-protein or protein-DNA interactions

To assess the ability of fluorescent partners to interact with Nanog, we developed a bead based coimmunoprecipitation assay termed coimmunoprecipitated bead imaging microscopy (CBIM). This uses confocal microscopy to directly image binding of a fluorescent molecule to a non-fluorescent partner protein immobilised on protein G-sepharose beads (Fig. S2A). Nuclear lysates from cells transfected with fluorescent protein expression constructs were used as a source of fluorescent proteins. CHO cells were used as the transfection target cell as these cells do not express components of the pluripotency gene regulatory network and therefore lack proteins that interact with the fluorescent protein fusions in a cell-specific manner. When beads coated with anti-FLAG antibody were incubated with a nuclear lysate from untransfected CHO cells, no fluorescence was detected on the beads by confocal microscopy (Fig. 1A). Likewise, when beads bound by anti-FLAG antibody were incubated with nuclear lysates from CHO cells transfected with separate constructs expressing (FLAG)_3_-Nanog and GFP, no fluorescence was apparent (Fig. 1B). However, incubation with lysate co-expressing (FLAG)_3_-Nanog and GFP-Nanog showed fluorescence localised on the bead surface, indicative of dimerization between (FLAG)_3_-Nanog bound to the anti-FLAG conjugated beads and GFP-Nanog from the CHO nuclear lysate (Fig. 1B). In contrast, when a GFP-Nanog fusion in which the tryptophan residues in the tryptophan repeat (WR; Nanog-W10A) was co-expressed alongside (FLAG)_3_-Nanog, incubation of CHO cell nuclear lysate with anti-FLAG-conjugated beads yielded no fluorescence (Fig. 1B). This is consistent with previous reports of a critical role for WR tryptophan residues in homodimerisation of Nanog (Mullin et al 2017, Mullin et al 2008, Wang et al 2008). In addition, incubation of beads with (FLAG)_3_-Nanog and GFP-Sox2, resulted in localisation of fluorescence to the bead surface (Fig. 1B). This is consistent with the known heterodimerisation of Nanog with Sox2 (Gagliardi et al 2013). We further assessed the heterodimerisation of Nanog and Sox2 by first immobilising (FLAG)_3_-Sox2 onto anti-FLAG/protein G-beads. Incubation with GFP-Nanog, but not GFP-Nanog-W10A resulted in fluorescent beads (Fig. 1C), consistent with a central role of WR tryptophans in mediating heterodimerisation of Nanog with Sox2 (Gagliardi et al 2013). We next assessed the utility of mCherry fusion proteins in CBIM (Fig. 1D). Binding of mCherry-NANOG or mCherry-SOX2 to anti-FLAG/proteinG-beads in the presence of (FLAG)_3_-Nanog established that interactions between partner proteins can be detected independent of the fluorescent fusion partner. These assays also showed that binding of Oct4 to Nanog could be detected by CBIM (Fig. 1D). Previously, Oct4 has been reported to co-immunoprecipitate with Nanog (Wang et al 2008). However, this result is not uncontentious, since Oct4 is not detectable by Nanog-affinity purification and mass spectrometry (Wang et al 2008) nor is Nanog present in the Oct4 interactome (van den Berg et al 2010). The current results confirm that Nanog and Oct4 can interact and suggest that CBIM may provide a sensitive means of detecting protein-protein interactions.

**Figure 1.**
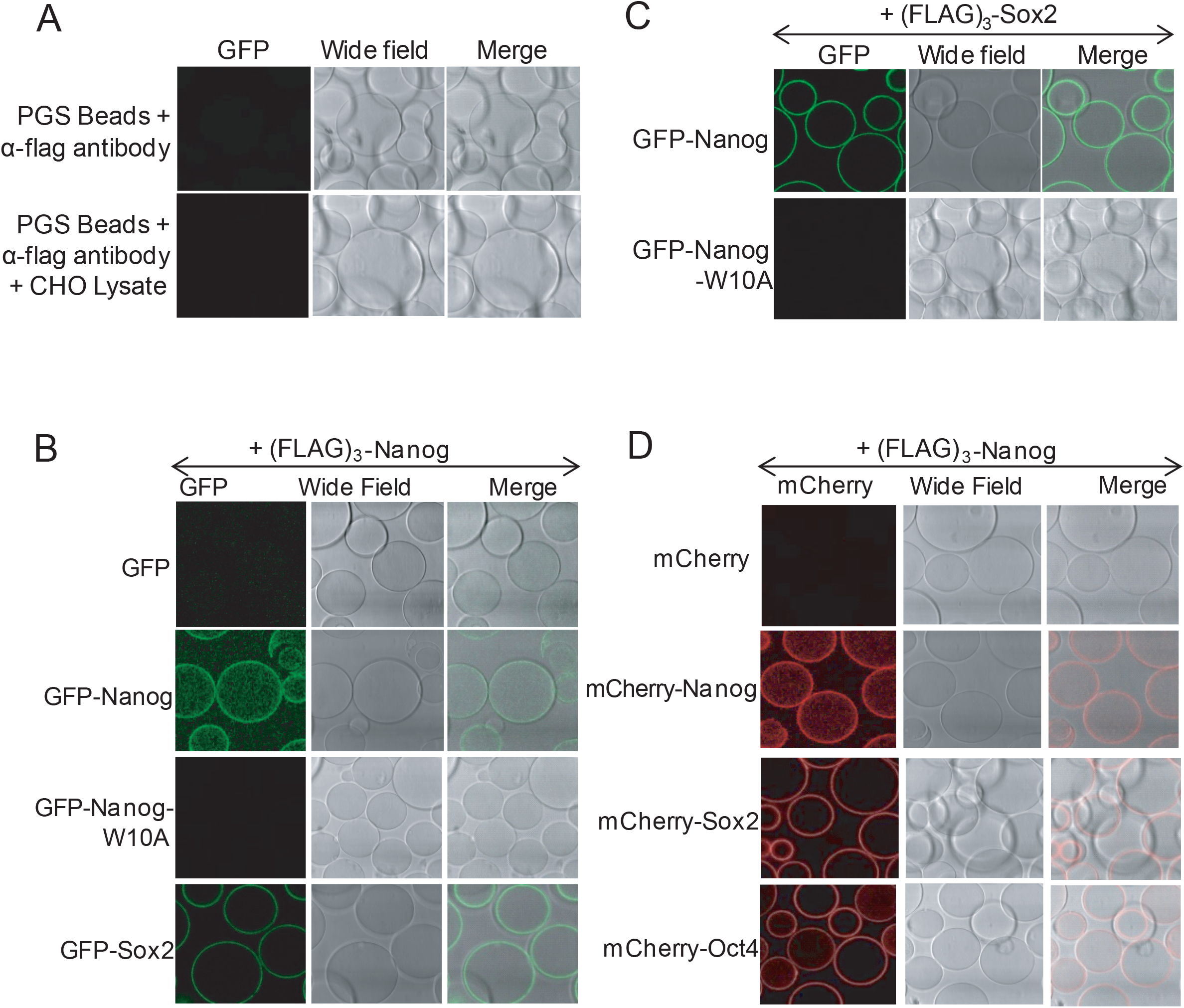
CBIM demonstrates Nanog binding to Sox2 and Oct4. **A**. Confocal images of protein-G beads incubated with antibody alone or antibody and CHO lysate. **B**. Confocal images of protein-G/anti-FLAG beads after CoIP from nuclear lysate of CHO cells co-transfected with (FLAG)_3_-Nanog and either GFP,GFP-Nanog, GFP-Nanog-W10A or GFP-Sox2. **C**. Confocal images of protein-G/anti-FLAG beads after CoIP from nuclear lysate of CHO cells co-transfected with (FLAG)_3_-Sox2 and either GFP-Nanog or GFP-Nanog-W10A. **D**. Confocal images of protein-G/anti-FLAG beads after CoIP from nuclear lysate of CHO cells co-transfected with (FLAG)_3_-Nanog and either mCherry, mCherry-Nanog, mCherry-Sox2 or mCherry-Oct4.

We next assessed the ability of CBIM to detect interactions between immobilised proteins and labelled DNA. (FLAG)_3_-Sox2 CHO cell lysate was first assessed by incubation ofanti-FLAG conjugated beads and a Cy5-labelled dsDNA containing the Oct/Sox motif from the *Nanog* gene (NSO) (Fig. S2B). This demonstrated fluorescence detectable on the bead surface (Fig.2A). This DNA interaction is specific, since Cy5-labelled dsDNA from the *Tcf3* gene that lacks an Oct/Sox motif was not retained on beads bound by Sox2 (Fig. 2A). We next assessed the ability of (FLAG)_3_-Oct4 to bind the same Cy5-labelled oligonucleotides to anti-FLAG/proteinG-beads and obtained similar results (Fig.2B). We then tested the ability of (FLAG)_3_-Nanog to localise DNA to the bead surface using the aforementioned Cy5-labelled dsDNA from the *Tcf3* gene (Fig. 2C). This oligonucleotide contains a Nanog binding site but lacks Oct4 or Sox2 binding sites (Jauch et al 2008). Incubating this oligonucleotide with nuclear lysate from CHO cells transfected with (FLAG)_3_-Nanog localised fluorescent DNA to the beads (Fig. 2C).

**Figure 2.**
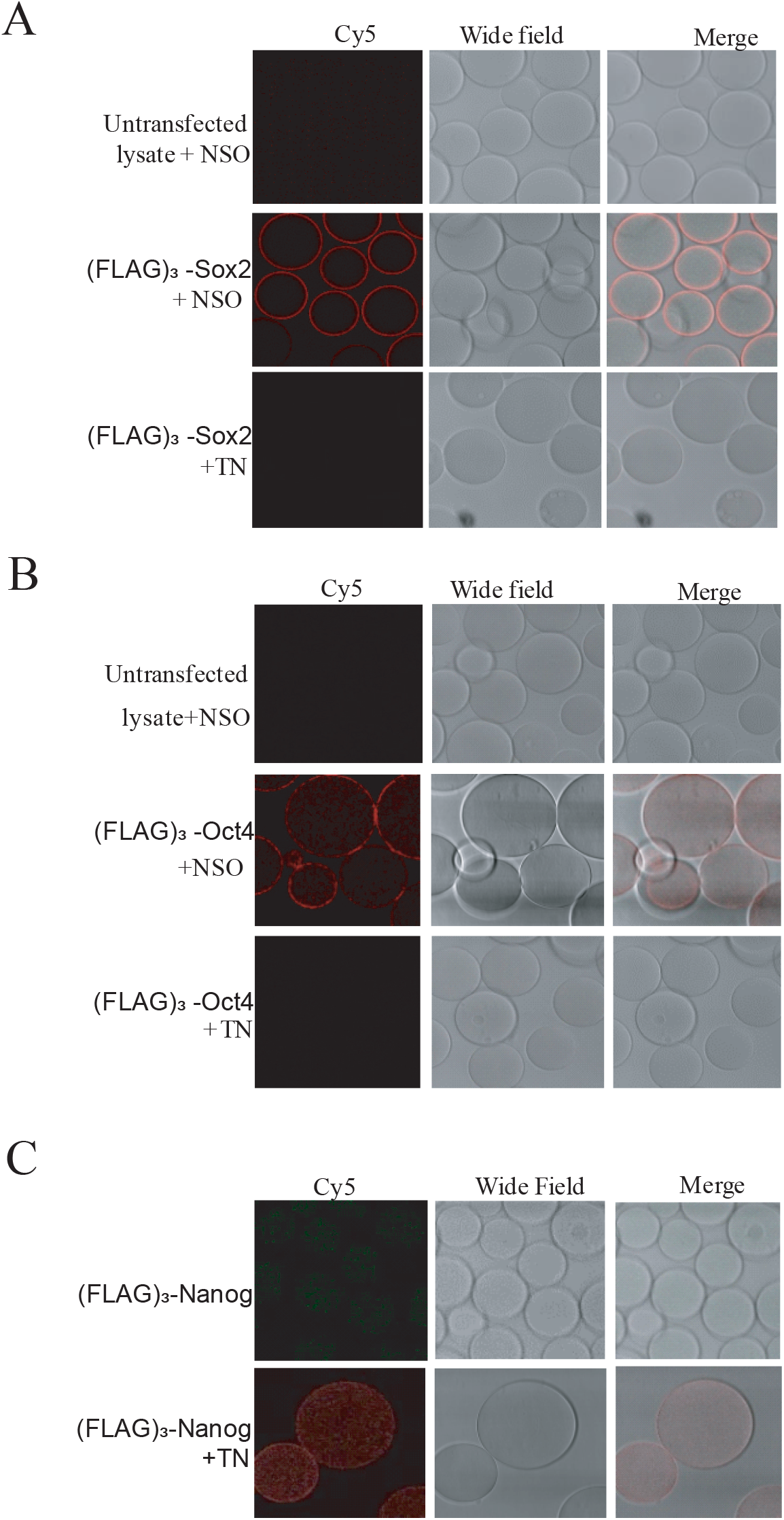
CBIM demonstrates DNA binding by TFs. **A**. Confocal images of protein-G/anti-FLAG beads incubated with a Cy5 labelled oligonucleotide encompassing the *Nanog* Sox/Oct (NSO) element and either CHO nuclear lysate from untransfected cells (upper panel) or (FLAG)_3_-Sox2 transfected cells (middle panel). In the lower panel a Cy5 labelled oligonucleotide encompassing the *Tcf3* specific Nanog (TN) element replaces the NSO sequence used in the middle panel. **B**. Confocal images of protein-G/anti-FLAG beads incubated with CHO nuclear lysate from untransfected cells (upper panel) or from cells transfected with (FLAG)_3_-Oct4 (middle and lower panels) together with the indicated DNA. **C**. Confocal images of protein-G/anti-FLAG beads incubated with CHO nuclear lysates transfected with (FLAG)_3_-Nanog, with or without Cy5 labelled TN DNA element. DNA sequences used are shown in Figure S2B.

### Nanog, but not Sox2 or Oct4 can form homo-multimers

To assess protein-protein complex formation at single molecule resolution, fluorescence correlation spectroscopy (FCS) was used. Nuclear lysates from CHO cells transfected with fluorescent protein fusions were examined by FCS to measure the photon <u>c</u>ount rate <u>p</u>er single <u>species</u> (CPS). Assessing the brightness of the molecules passing through the detection beam enables extraction of information about their multimeric state. These analyses show that the molecular brightness of mCherry and mCherry-Sox2 are similar, suggesting Sox2 does not form multimers (Fig. 3A). In contrast, mCherry-Nanog is brighter than mCherry, with the mCherry-Nanog/mCherry brightness ratio close to 2.0 (Fig. 3A). These results suggest that Nanog is in equilibrium between dimeric and monomeric states. FCS analysis of GFP fusions also confirmed that SOX2 is monomeric and that NANOG exists in equilibrium between monomeric and dimeric states. Moreover, mutation of the WR tryptophan residues eliminates multimerization capacity, consistent with Nanog-W10A existing as a monomer (Mullin et al 2008) (Fig. 3B).

**Figure 3.**
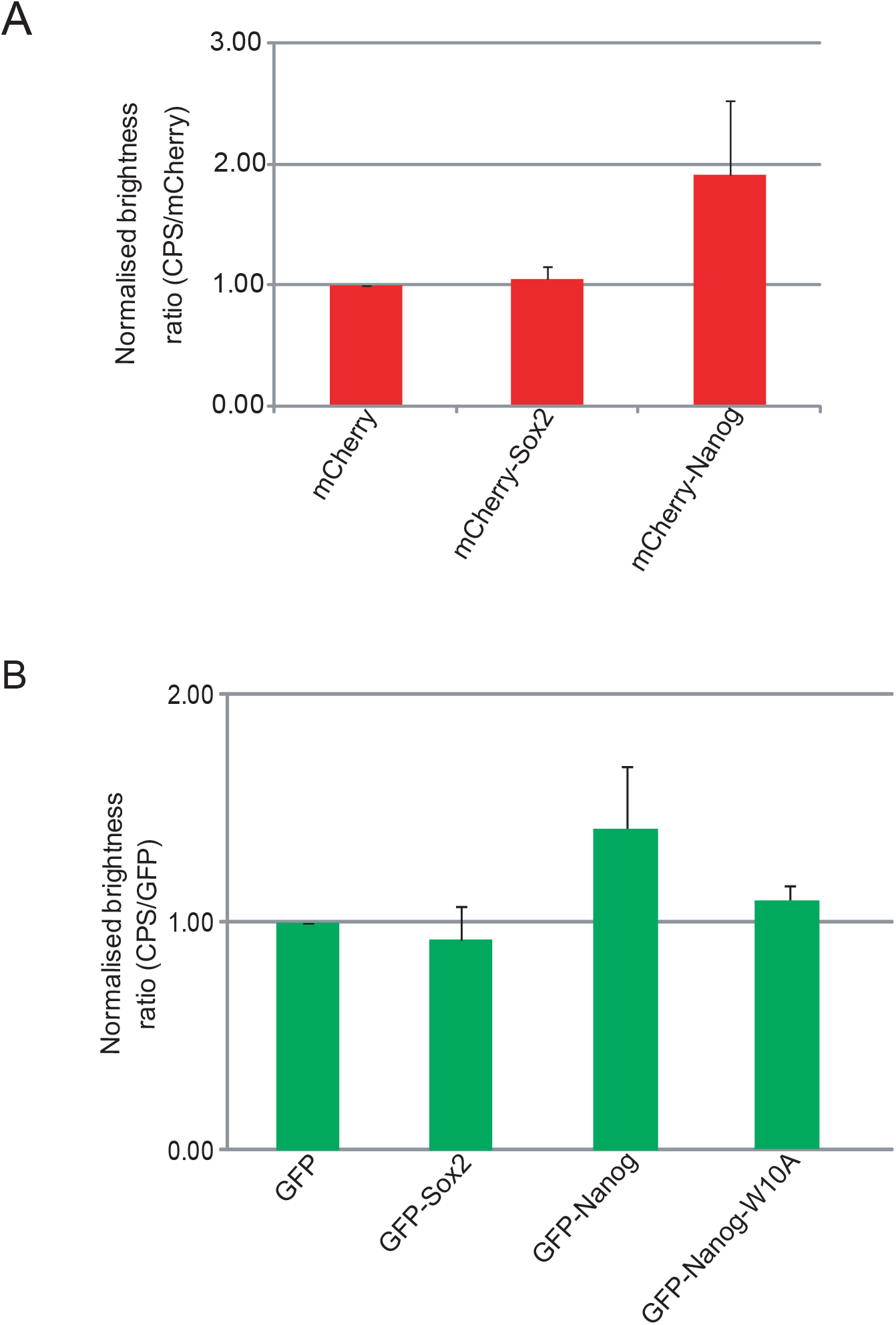
FCS analysis of transcription factors in nuclear lysates. **A**. Count rate per species (CPS) values for mCherry, mCherry-Sox2 and mCherry-Nanog measured by FCS. **B**. CPS of GFP, GFP-Sox2, GFP-Nanog, and GFP-Nanog-W10A were measured by FCS. For all experiments, n = 12, mean ± SD.

### Comparative assessment of the binding strengths of Nanog with partner proteins

The previous results suggest that Nanog is in equilibrium between monomers and dimers. To further assess the properties of Nanog dimers, we performed fluorescence cross-correlation spectroscopy (FCCS) experiments using mCherry and GFP fusions to Nanog, Sox2 and Oct4 (Fig. 4). By analysing the cross correlation between fluorescent protein signals originating from distinct partners in a dimer, we were able to assess the complex %, a parameter indicating the proportion of complex present in equilibrium and hence the affinity of binding (Table 1). Analysing CHO nuclear lysates expressing GFP and mCherry (Fig. 4A) or a tandem GFP-mCherry (Fig. 4A) provided the baseline and maximum for fluorescent proteins that are either separate, or always in complex. The calculated complex % for the negative control was 2.5 ± 1.9 and for the positive control 46.2 ± 6.7 (Fig. 4A, Table 1). FCCS of lysate expressing mCherry-Nanog and GFP-Nanog gave a complex % of 19.8 ± 2.9 (Fig. 4C,Table 1). Similar analysis of lysates expressing mCherry-Nanog with either GFP-Sox2 or GFP-Oct4 gave complex percentages of 26.1 ± 7.7 and 12.5 ± 6.7, respectively (Fig. 4C,Table 1). In contrast, FCCS analysis of GFP-Nanog-W10A in combination with mCherry-Nanog, mCherry-Sox2 or mCherry-Oct4 gave complex percent values indistinguishable from the negative control (Fig. 4C, Table 1). These results suggest an order of binding strength of Nanog for partners of Sox2>Nanog> Oct4.

**Table 1:** Complex% values for different sets of *in-vitro* FCCS experiments from Figure 4.

| Interacting partners | Complex % |
| --- | --- |
| **Interaction between two proteins** |  |
| GFP-Oct4 + mCherry-Sox2 | 3.9 ± 3.1 |
| GFP-Oct4 + mCherry-Nanog | 12.5 ± 6.7 |
| GFP-Sox2 + mCherry-Nanog | 26.1 ± 7.7 |
| GFP-Nanog+ mCherry-Nanog | 19.8 ± 2.9 |
| **Interaction between three proteins** |  |
| GFP-Oct4 + mCherry-Sox2 + (Flag)<sub>3</sub>-Nanog | 3.0 ± 2.7 |
| GFP-Oct4 + mCherry-Nanog + (Flag)<sub>3</sub>-Sox2 | 5.0 ± 3.5 |
| **Positive, negative and mutation controls** |  |
| GFP-mCherry (tandem) | 46.2 ± 6.7 |
| GFP + mCherry | 2.5 ± 1.9 |
| mCherry-Nanog + GFP-Nanog-W10A | 1.7 ± 1.4 |
| mCherry-Oct4 + GFP-Nanog-W10A | 3.6 ± 2.5 |
| mCherry-Sox2 + GFP-Nanog-W10A | 2.8 ± 1.7 |

**Figure 4.**
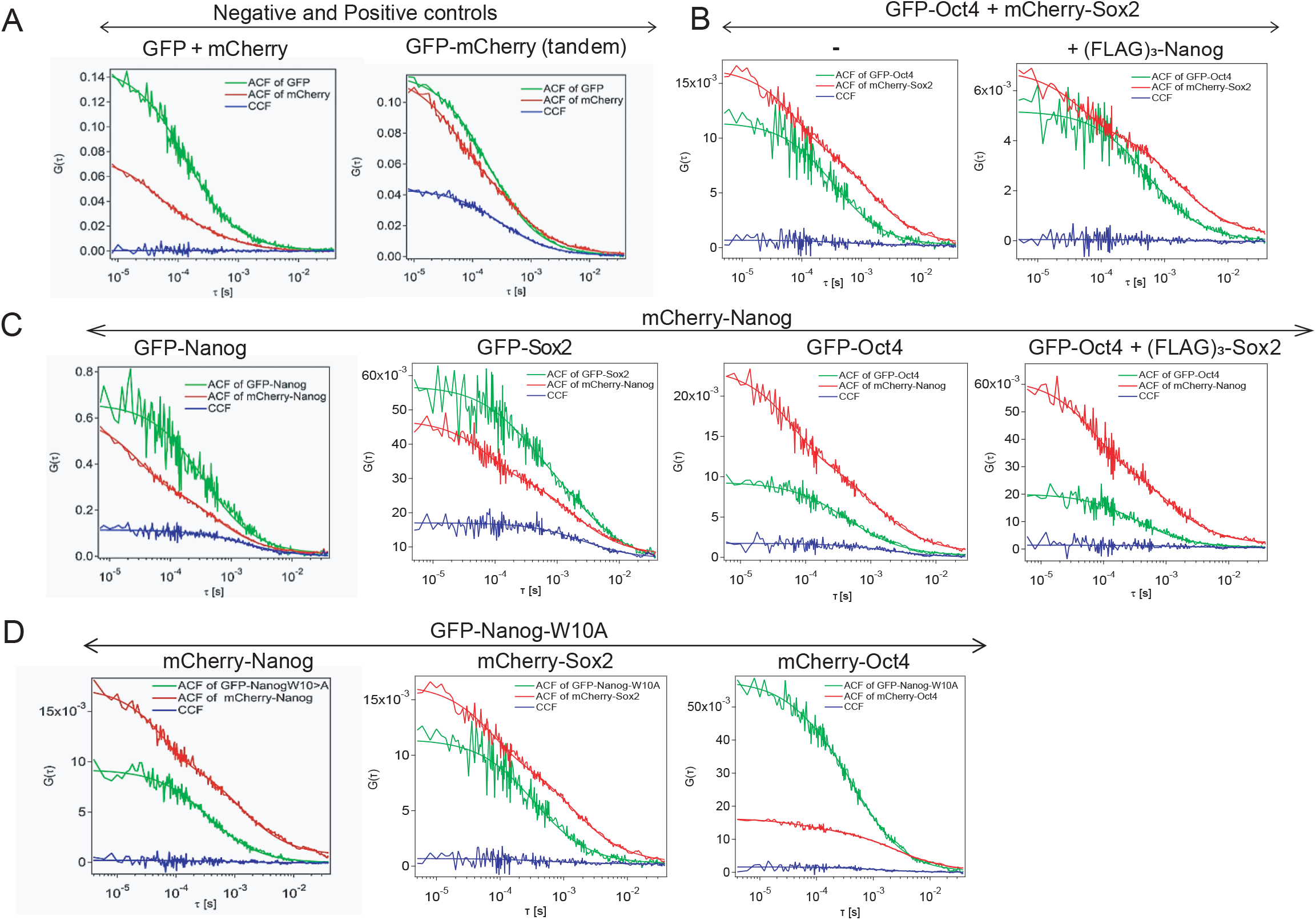
FCCS analysis demonstrates interaction between Nanog, Sox 2 and Oct4. **A**. Autocorrelation function and cross-correlation function curves from FCCS assay of nuclear lysates containing co**-**transfected GFP and mCherry (left) or containing a tandem GFP-mCherry protein (right). **B**. Autocorrelation function (ACF) and cross-correlation function (CCF) curves from FCCS with nuclear lysate from a co-transfection of GFP-Oct4 and mCherry-Sox2 in the presence and absence of (FLAG)_3_-Nanog. **C**. Autocorrelation function (ACF) and cross-correlation function (CCF) curves from FCCS assay of nuclear lysates containing mCherry-Nanog co-transfected with either GFP-Nanog, GFP-Sox2, GFP-Oct4 or both of GFP-Oct4 and (FLAG)_3_-Sox2. **D**. Autocorrelation function and cross-correlation function curves from FCCS assay of nuclear lysate containing GFP-Nanog-W10A co-transfected with mCherry-Nanog, mCherry-Sox2 or mCherry-Oct4. For all experiments, n = 12, mean ± SD. The percentage complex formation calculated from the above assays are shown in Table 1.

### Nanog diffusion dynamics decrease in the presence of either Sox2 or Oct4

To obtain insight into the dynamics of the interaction of Nanog with Sox2 or Oct4 we used fluorescence correlation spectroscopy (FCS) to measure the diffusion coefficient of GFP-Nanog alone, or in the presence of either (FLAG)_3_-Sox2 or (FLAG)_3_-Oct4 (Fig. 5, Fig. S3). This showed that the diffusion coefficient of GFP-Nanog decreases to a similar extent in the presence of either Oct4 or Sox2. As Sox2 and Oct4 have a similar molecular weight, this suggests that the Nanog complexes they form may have similar stoichiometries. Moreover, as the diffusion coefficient of GFP-Nanog is further reduced by the presence of both Sox2 and Oct4, this suggests that Nanog, Sox2 and Oct4 can form a ternary complex in solution. Interestingly, GFP-Nanog-W10A has a higher diffusion coefficient than GFP-Nanog, consistent with monomer formation by GFP-Nanog-W10A (Mullin et al 2008). Moreover, as the diffusion coefficient of GFP-Nanog-W10A is unaffected by addition of Sox2 or Oct4, this suggests that the WR tryptophan residues in NANOG are required for interaction with Oct4, as has been shown for SOX2 (Gagliardi et al 2013).

**Figure 5.**
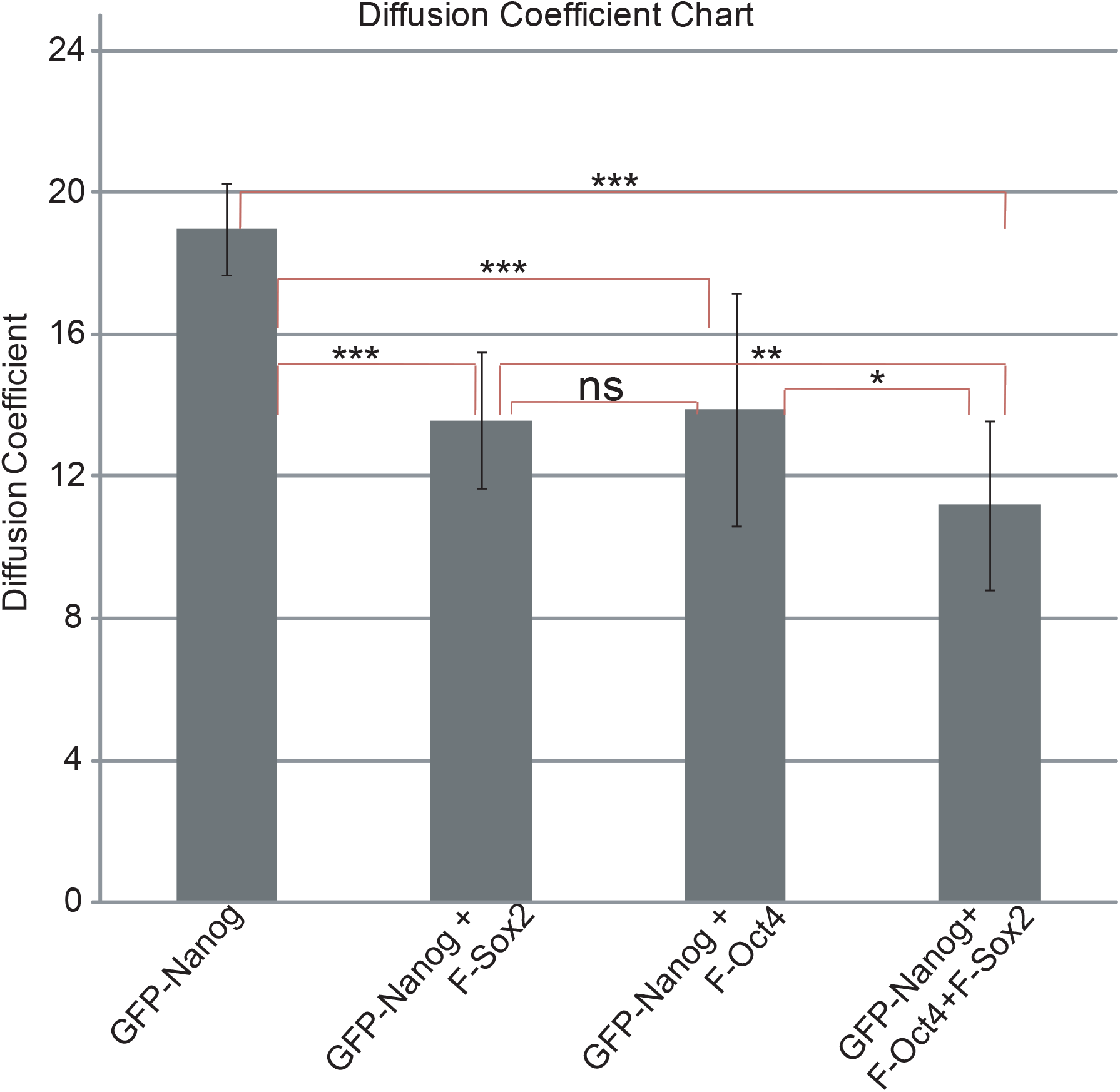
The diffusion dynamics of Nanog change in the presence of co-factors. Diffusion coefficients of GFP-Nanog and GFP-Nanog in the presence of either (FLAG)_3_-Sox2, (FLAG)_3_-Oct4 or both of (FLAG)_3_-Sox2 and (FLAG)_3_-Oct4 were measured by FCS. F-Sox2 and F-Oct4 stand for (FLAG)_3_-Sox2 and (FLAG)_3_-Oct4 respectively. Statistical significance was measured using an unpaired t-test. p values are indicated (<0.0002, ***; 0.0233, **, 0.0508, *; ns, not significantly different). For all experiments n = 12, mean ± SD.

### Competitive interactions between TF partners

We next extended the FCCS analysis to assess the Oct4-Sox2 interaction using nuclear lysates from CHO cells co-transfected with GFP-Oct4 and mCherry-Sox2 (Fig. 4B). This showed that Sox2 does not bind to Oct4 in solution (Complex % = 3.9 ± 3.1) (Table 1) (Chen et al 2012) and that addition of Nanog did not affect the extent of the interaction between Oct4 and Sox2 (Fig. 4B, Table 1). Therefore, Nanog neither assists nor impairs the interaction between Oct4 and Sox2 in solution.

In contrast, analysis of the interaction between GFP-Oct4 and mCherry-Nanog showed that the complex % was reduced from 12.5 (±6.7) to 5.0 (±3.5) upon addition of (FLAG)_3_-Sox2 (Fig. 4C, Table 1). There are at least two possible interpretations of this data. Firstly, Sox2 might bind to Oct4 and thereby reduce the interaction between Oct4 and Nanog. However, as Sox2 binding to Oct4 is DNA-dependent (Chen et al 2012), this seems unlikely. An alternative is that Sox2 and Oct4 may compete for binding to Nanog and that Sox2 might preferentially bind to Nanog. This possibility is supported by the observation that both Sox2 and Oct4 reduce the diffusion co-efficient of GFP-Nanog but neither affect the diffusion of GFP-Nanog-W10A, implying that Oct4, like Sox2 binds Nanog via the WR tryptophan residues (Fig. 5).

### Analysis of Nanog dimer formation *in vivo*

The preceding analyses indicate that Nanog can interact with partners on beads and in solution. To investigate whether similar interactions may occur *in vivo*, we assessed live cells, initially using FCS to analyse fluorescent protein brightness in CHO cells transfected with either GFP or GFP-Nanog (Fig. 6). Similar to the situation *in vitro*, the brightness of GFP-Nanog was higher than that of GFP, with a GFP-Nanog/GFP ratio of approximately 1.5 (Fig. 6B). These results are consistent with the idea that Nanog monomers and multimers are in equilibrium within cells.

**Figure 6.**
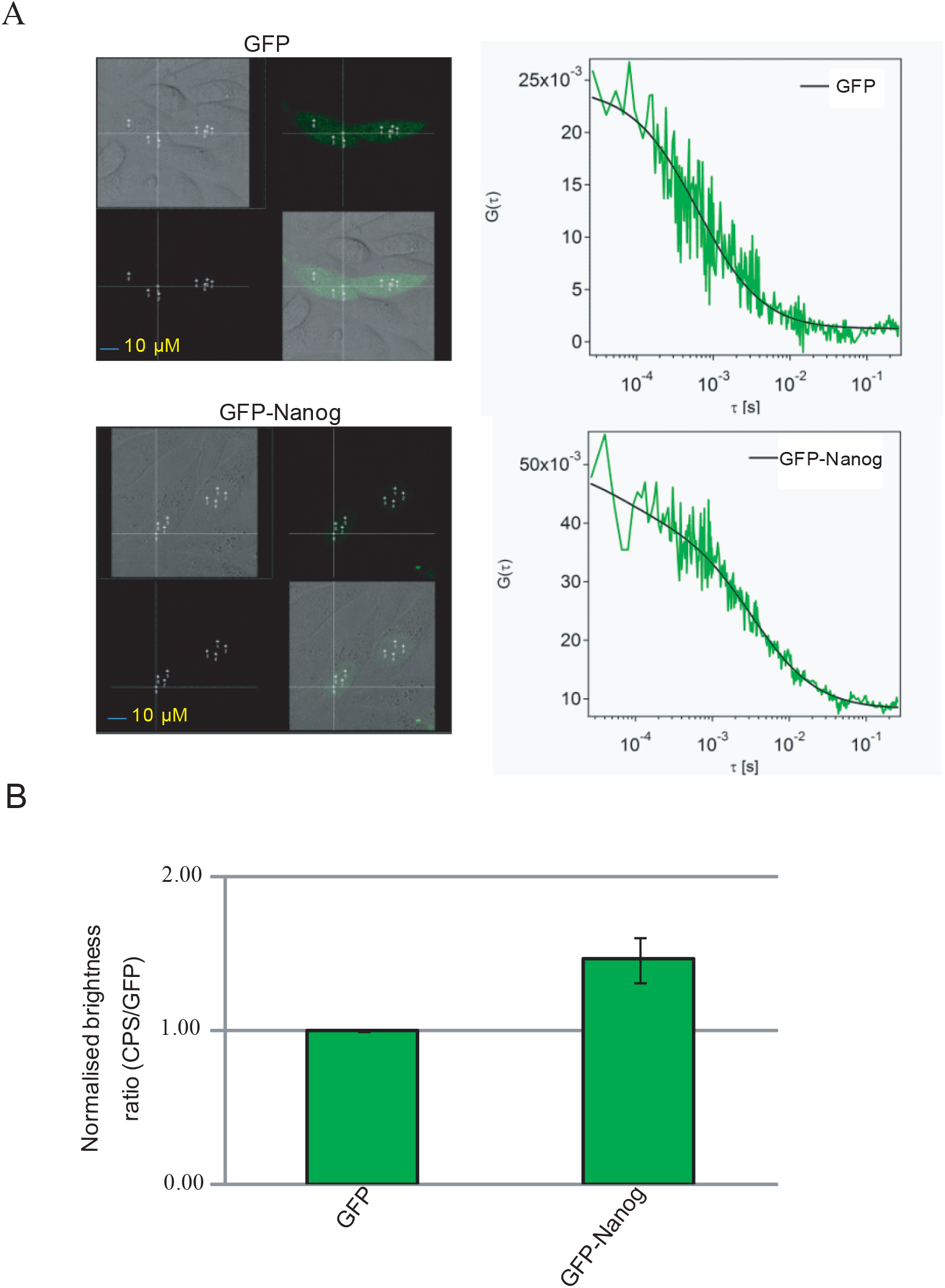
FCS analysis of transcription factors in live cells. **A**. Confocal images (left) and best-fit autocorrelation function (ACF) curves generated from FCS assay of cells expressing GFP (top) or GFP-Nanog (bottom). Scale bar = 10µm. **B**. Normalized count rate per species (CPS) values for GFP and GFP-Nanog. For all experiments, n = 15, mean ± SD.

The preceding FCS analysis is consistent with the existence of Nanog dimers *in vivo*. However, they do not provide information about the proportion of molecules present in monomeric or multimeric forms *in vivo*. To assess this, we used fluorescence cross-correlation spectroscopy to analyse live CHO cells co-transfected with Nanog-GFP and mCherry-Nanog (Fig. 7, Fig. S4). In cells expressing GFP and mCherry as a negative control, the baseline complex %, a measure of binding strength, was 3.7 ± 2.6 whereas that for the GFP-mCherry fusion, the positive control, complex % was 47.4 ± 9.2 (Fig. 7A). These values then served as the lower and upper limits of the complex % for experiments in which mCherry-Nanog was expressed in CHO cells alongside GFP-Nanog or GFP-Nanog-W10A. FCCS data indicate that the cross correlation between GFP-Nanog and mCherry-Nanog, indicated a complex % of 16.8 ± 7.2 (Fig. 7B, C), a value close to that obtained from *in vitro* analyses (Table 1). In contrast, the level of complex formed by mCherry-Nanog and GFP-Nanog-W10A was similar to that of mCherry + GFP (Fig. 7A, B, C), indicating no interaction between mCherry-Nanog and GFP-Nanog-W10A. These results indicate that Nanog exists in cells in an equilibrium between monomeric and multimeric forms.

**Figure 7.**
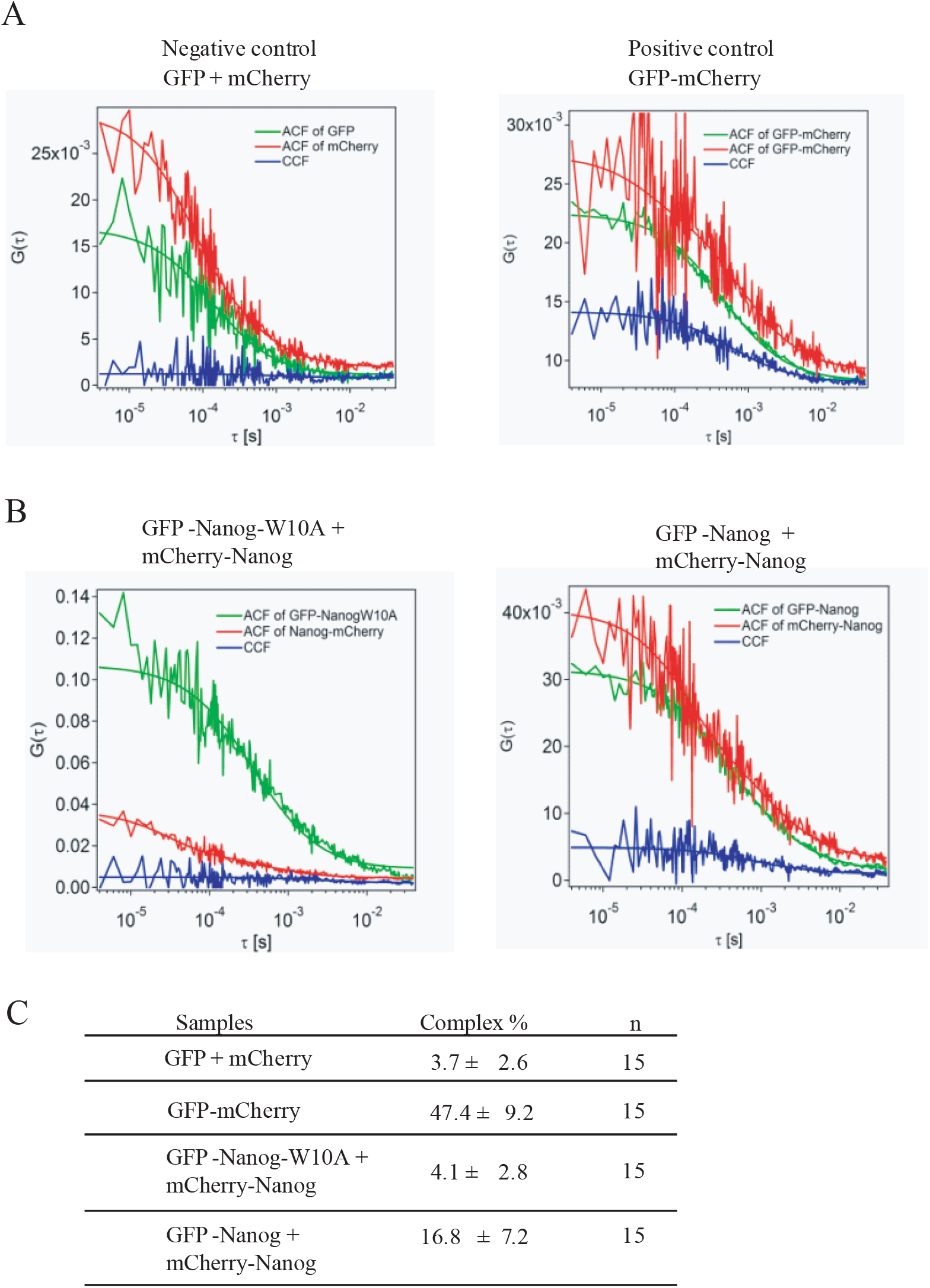
FCCS analysis of transcription factors in live cells. **A**. Autocorrelation functions (ACF) and cross-correlation functions (CCF) from FCCS assay with live CHO cells co-transfected with GFP and mCherry (left) or with tandem GFP-mCherry (right). **B**. Autocorrelation functions and cross-correlation functions from live CHO cells co-transfected with mCherry-Nanog and either GFP-Nanog-W10A (left) orGFP-Nanog (right). **C**. Percentage complex for species in A and B. For all experiments, cell number = 15, mean ± SD).

## DISCUSSION

In this study, we assess the molecular dynamics of Nanog in solution at single molecule sensitivity. It has previously been shown that Nanog homodimerises and heterodimerises with Sox2 to form DNA independent protein-protein complexes (Gagliardi et al 2013, Mullin et al 2008, Wang et al 2008). However, the quantitative details of Nanog protein-protein complex formation have only partly been addressed. Our analysis of the diffusion dynamics of Nanog fluorescent protein fusions confirms that two Nanog molecules participate in Nanog homo-multimerisation and that dimers remain in equilibrium with monomers, consistent with previous analytical ultracentrifugation studies (Mullin et al 2008). Importantly, our FCCS analysis establishes the existence of Nanog homomultimers in cells.

FCS analysis shows that the capacity of Nanog for homo-multimerisation requires tryptophan residues within the Nanog WR of both mouse and human Nanog (Choi et al 2022, Mullin et al 2008, Wang et al 2008). This capacity for homo-multimerisation is not shared with Sox2. This conclusion is substantiated by analysis of diffusion coefficients of fluorescent protein which indicates that GFP-Nanog forms larger complexes than GFP-Nanog-W10A. Morevover, while the diffusion of GFP-Nanog is reduced by addition of either Oct4 or Sox2, this does not occur upon addition of either Sox2 or Oct4 to GFP-Nanog-W10A. This indicates that Oct4 and Sox2 each interact with Nanog via WR tryptophan residues.

In addition, by inclusion of a non-fluorescent partner with a FCCS pair it is possible to determine the effect of the addition on dimerization. Thus, addition of Sox2 alongside GFP-Oct4 and mCherry-Nanog reduced the amount of the Oct4/Nanog complex, likely through a competition between Oct4 and Sox2 for binding to Nanog.

We describe a new confocal microscopy method to detect binding of a fluorescently labelled protein or DNA to a target protein (coimmunoprecipitated bead imaging microscopy [CBIM]). Using CBIM allowed us to establish that Nanog binds Oct4, a previously contentious issue (Gagliardi et al 2013, van den Berg et al 2010, Wang et al 2008). The high local concentration of binding partners that can be achieved on the bead surface may make CBIM a sensitive method for detecting low affinity interactions. This idea is supported by our FCCS analysis of the capacity of Nanog to form homodimers, or heterodimers with Sox2 or Oct4. This data indicates that the affinity of Nanog for binding partners is in the order Sox2 > Nanog > Oct4.

## Supporting information

All supplementary Figures

## ACKNOWLEDGEMENTS

Research in the IC lab was supported by the Medical Research Council of the UK.

## AUTHOR CONTRIBUTIONS

T.K.M. did experiments and wrote the manuscript, D. K. helped T.K.M with fluorescent imaging experiments, J. M. performed diffusion study of Nanog, D.C. performed fusion Nanog functional tests, N.M. provided reagents, advice and assisted in manuscript preparation and I.C. supervised the project and wrote the manuscript.

## ADDITIONAL INFORMATION

Please see the **Supplementary Section** for further information.

## FIGURE LEGENDS

**Supplementary Figure 1. Functional test of Nanog fusion proteins in self-renewal assay**. E14/T ES cells were transfected with plasmids encoding Nanog fusion proteins and colonies stained with alkaline phosphatase 10 days post-transfectionin the absence of LIF. Undifferentiated (ES), mixed and differentiated colonies were counted. n = 3, mean ± SD.

**Supplementary Figure 2. CBIM diagram. A**. Protein G beads are coated with anti-FLAG antibody and used to immunoprecipitateproteins or complexes of interest. The protein of interest or a binding partner is fused to a fluorescent protein allowing interactions to be visualized by confocal microscopy. The assay can also be used to visualize the interaction between protein and a fluorescently tagged DNA target sequence. **B**. DNA sequences used in Figure 2.

**Supplementary Figure 3. The diffusion dynamics of Nanog change in the presence of co-factors**. Diffusion coefficients of GFP, GFP-Nanog-W10A, and GFP-Nanog were measured by FCS, in the presence or absence of (FLAG)_3_-Sox2 or (FLAG)_3_-Oct4 as indicated. F-Sox2 and F-Oct4 stand for (FLAG)_3_-Sox2 and (FLAG)_3_-Oct4 respectively. Statistical significance was measured using an unpaired t-test. For all experiments n = 12, mean ± SD.

**Supplementary Figure 4. Confocal images of live cells used in FCCS analysis**. (**A-B**). Representative confocal images for positive (tandem GFP-mCherry) (A) and negative (co-transfection of GFP and mCherry) controls (B). Scale bar is 3.5 µm. (**C-D**). Representative confocal images for CHO cells co-transfected cells with GFP-Nanog-W10A and mCherry-Nanog (**C**) and CHO cells co-transfected with GFP-Nanog and mCherry-Nanog (**D**). The scale bar is 3.5 µm. The intersection of the two lines refers to the position where FCCS was measured.

