## Supplementary material for "Analysis of pluripotency transcription factor interactions reveals preferential binding of NANOG to SOX2 rather than NANOG or OCT4": All supplementary Figures

Mistri et al
Supplementary Figure 1

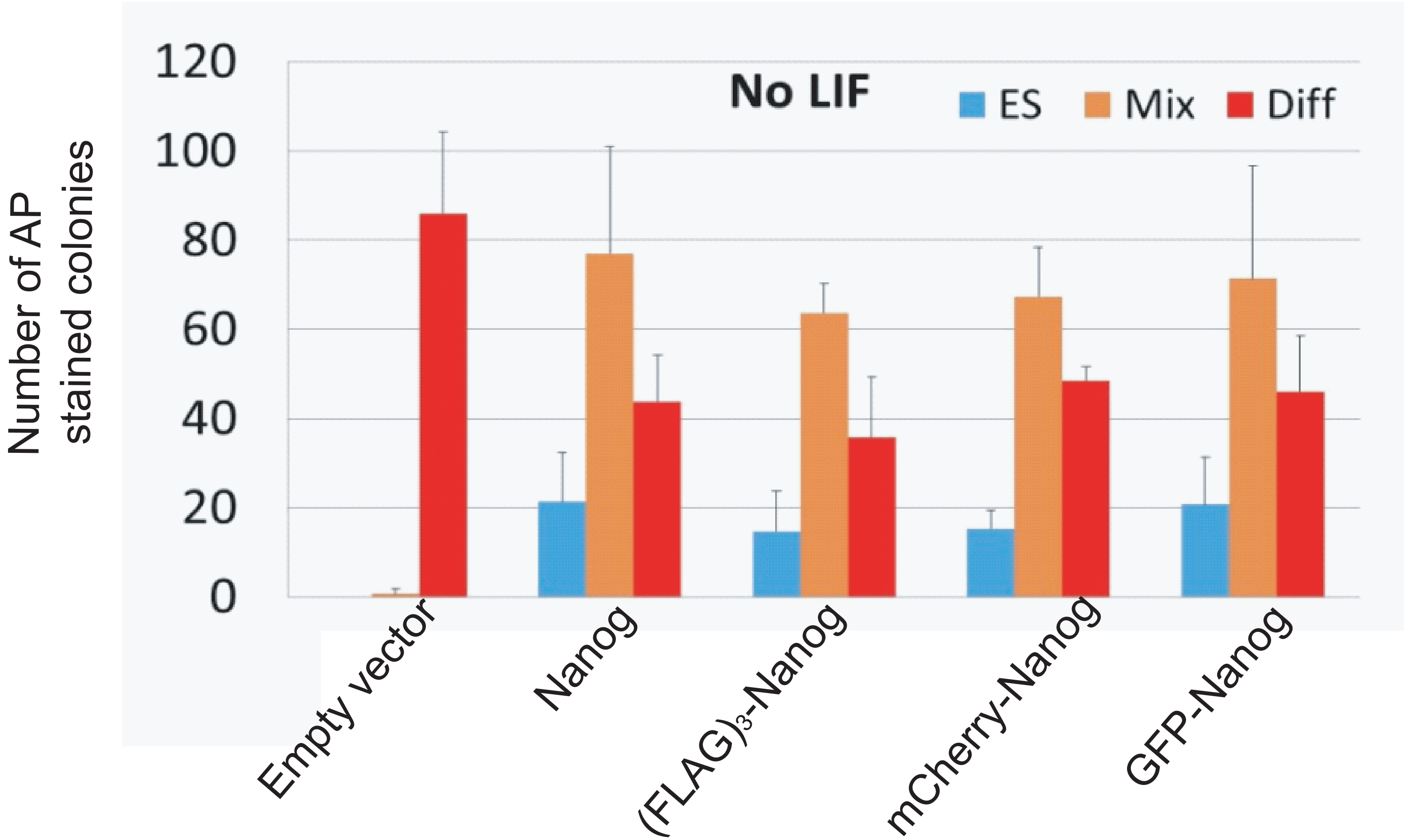

A

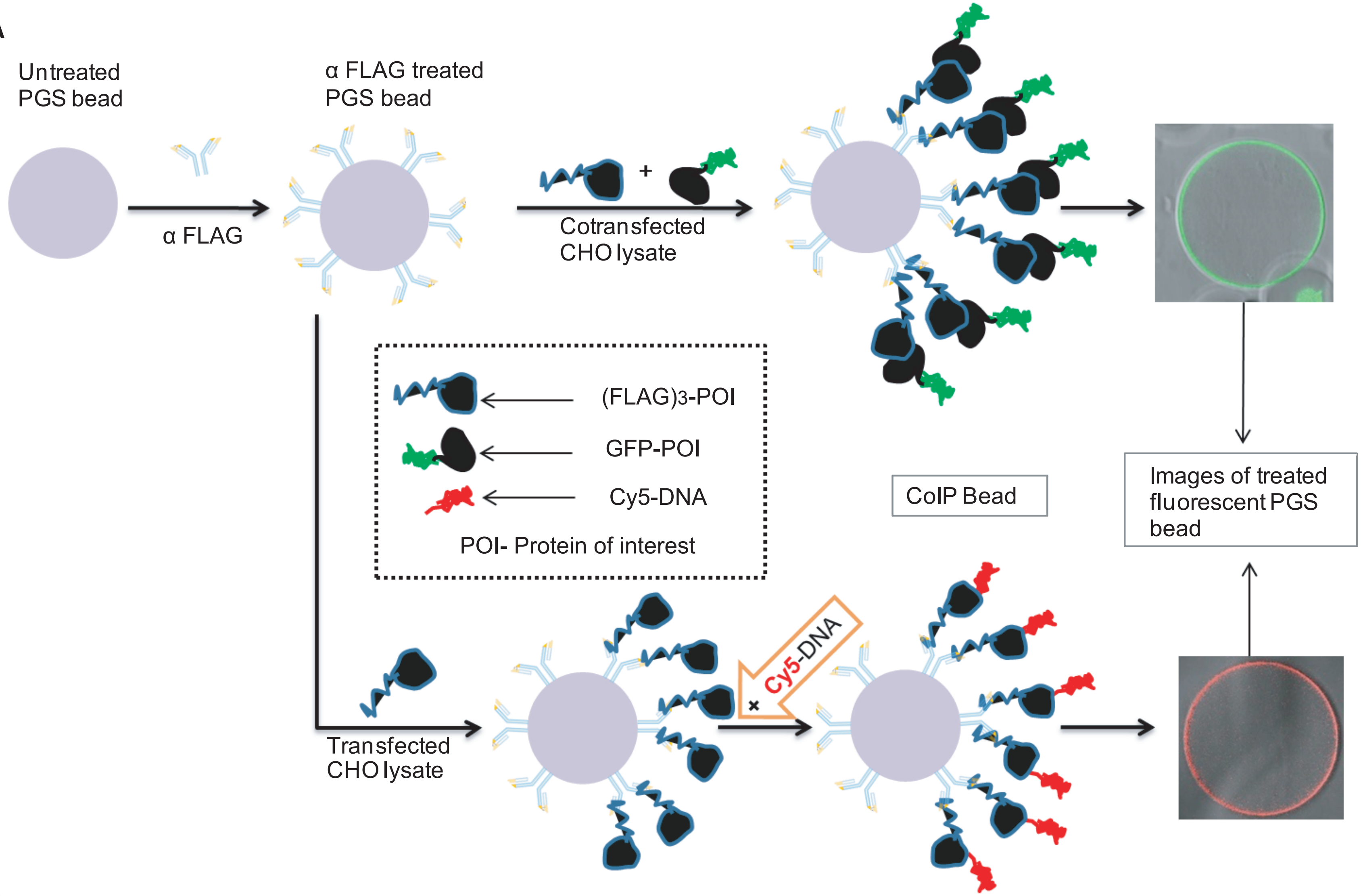


B

| Name of DNA motif | DNA Sequence |
|---|---|
| Nanog Sox/Oct *cis* motif (**NOS**) | 5'-Cy5-GGACATTGTAATGCAAAAGAA -3' |
| Tcf3 Nanog *cis* motif (**TN**) | 5'-Cy5-AACCTGTTAATGGGAGCG -3' |

Mistri et al
Supplementary Figure 3

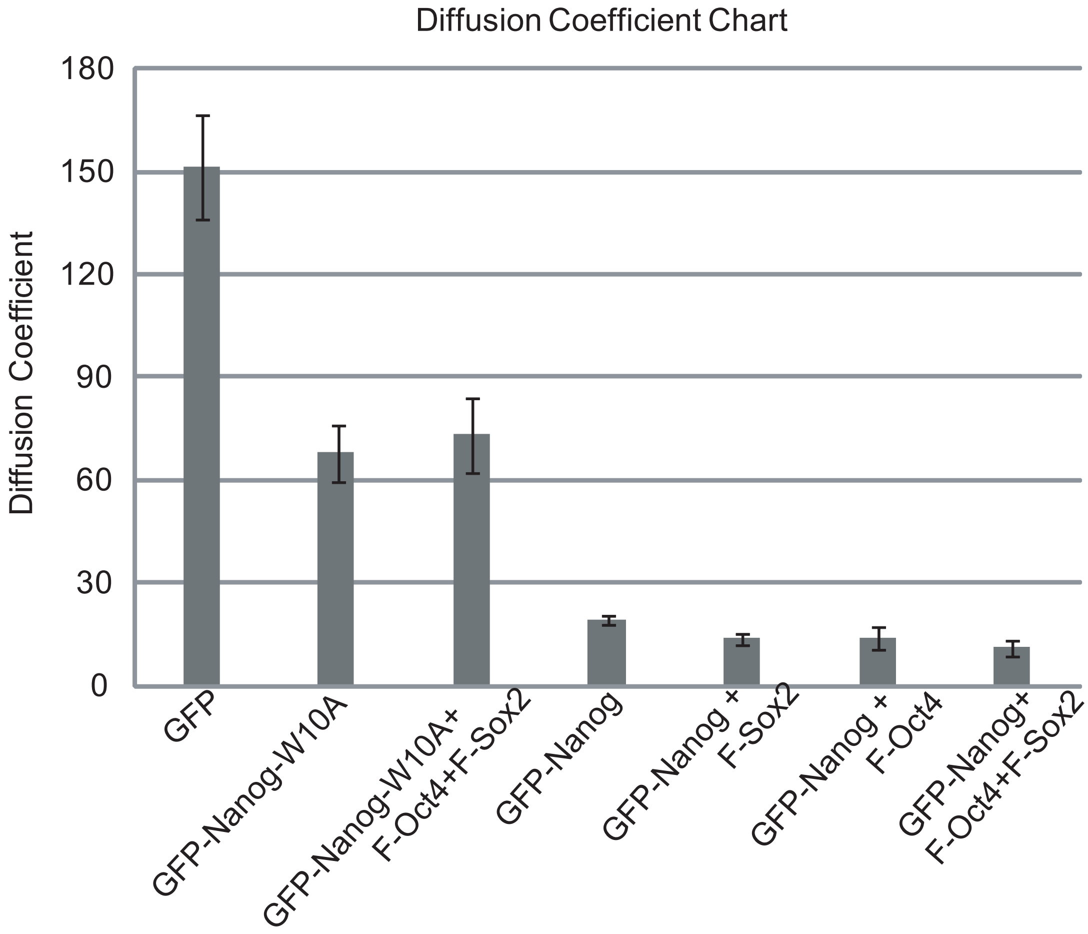

Mistri et al
Supplementary Figure 4

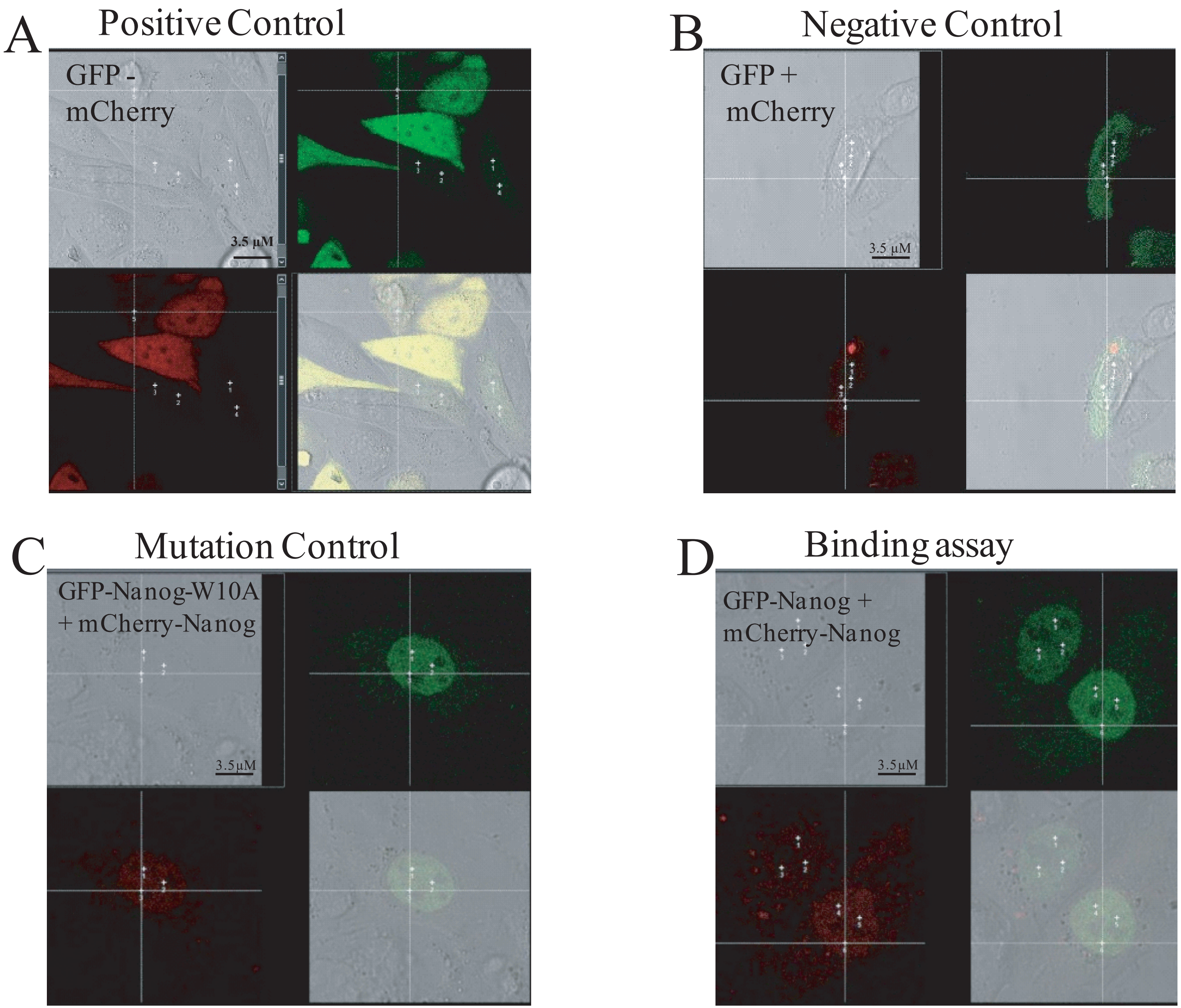
